# Unconstrained Precision Mitochondrial Genome Editing with αDdCBEs

**DOI:** 10.1101/2024.05.13.593977

**Authors:** Santiago R. Castillo, Brandon W. Simone, Karl J. Clark, Patricia Devaux, Stephen C. Ekker

## Abstract

DddA-derived cytosine base editors (DdCBEs) enable the targeted introduction of C•G-to-T•A conversions in mitochondrial DNA (mtDNA). DdCBEs are often deployed as pairs, with each arm comprised of a transcription activator-like effector (TALE), a split double-stranded DNA deaminase half, and a uracil glycosylase inhibitor. This pioneering technology has helped improve our understanding of cellular processes involving mtDNA and has paved the way for the development of models and therapies for genetic disorders caused by pathogenic mtDNA variants. Nonetheless, given the intrinsic properties of TALE proteins, several target sites in human mtDNA remain out of reach to DdCBEs and other TALE-based technologies. Specifically, due to the conventional requirement for a thymine immediately upstream of the TALE target sequences (i.e., the 5’-T constraint), over 150 loci in the human mitochondrial genome are presumed to be inaccessible to DdCBEs. Previous attempts at circumventing this constraint, either by developing monomeric DdCBEs or utilizing DNA-binding domains alternative to TALEs, have resulted in suboptimal specificity profiles with reduced therapeutic potential. Here, aiming to challenge and elucidate the relevance of the 5’-T constraint in the context of DdCBE-mediated mtDNA editing, and to expand the range of motifs that are editable by this technology, we generated αDdCBEs that contain modified TALE proteins engineered to recognize all 5’ bases. Notably, 5’-T-noncompliant, canonical DdCBEs efficiently edited mtDNA at diverse loci. However, DdCBEs were frequently outperformed by αDdCBEs, which consistently displayed significant improvements in activity and specificity, regardless of the 5’-most bases of their TALE binding sites. Furthermore, we showed that αDdCBEs are compatible with DddA_tox_ and its derivatives DddA6, and DddA11, and we validated TALE shifting with αDdCBEs as an effective approach to optimize base editing outcomes at a single target site. Overall, αDdCBEs enable efficient, specific, and unconstrained mitochondrial base editing.

## INTRODUCTION

Mitochondria are semi-autonomous organelles with a central role in energy metabolism that contain a circular and multicopy genome, mitochondrial DNA (mtDNA), which in humans encodes 37 genes critical for oxidative phosphorylation.^1–7^ Pathogenic mtDNA variants are prevalent in ∼1 in 5,000 people and are causal in currently incurable metabolic disorders.^8–12^ Mitochondrial base editing has recently emerged as a potential therapeutic approach for these mtDNA-based diseases.^13–16^ Notably, given the challenging nature of exogenous RNA import into mitochondria, the effective manipulation of mtDNA is enabled by all-protein systems.^16–20^ In light of their rapid and accessible engineering, transcription activator-like effectors (TALEs) are the most commonly used DNA-binding domains in most current mitochondrial base editing technologies.^16,21–36^

DddA-derived cytosine base editors (DdCBEs) consisting of pairs of mitochondrially targeted fusion proteins composed of a TALE, one half of a split dsDNA deaminase toxin (DddA_tox_), and a uracil glycosylase inhibitor, represent the most extensively developed tools for mtDNA editing.^16,30–34^ Based on prior work on TALE-DNA interactions in native contexts and in gene targeting applications in the nuclear compartment,^23–27,37–47^ the target flexibility of DdCBEs on mtDNA is presumed to be constrained by the requirement of a 5’ thymine (5’-T) in their TALE binding sites, restricting over 150 loci in the human mitochondrial genome.^13,48^ This design guideline stems from the specific interaction between the highly conserved N-terminal domain (NTD) of wild-type TALEs and the 5’-T of their target sequences.^37–40^ Despite being generally followed, the significance of this constraint for the design of effective DdCBEs has yet to be characterized.^13,16,30–34^

Here, we sought to elucidate the relevance of the TALE 5’-T rule for DdCBE-mediated mitochondrial base editing, as well as expand the targeting scope and design flexibility of this technology. To this end, building on our recently established system for the assembly of TALE-guided deaminases,^49–51^ we generated αDdCBEs that contain NT-αN, a previously developed TALE NTD engineered to circumvent the 5’-T rule.^52^ Subsequently, we conducted direct comparisons between DdCBEs and αDdCBEs in six mitochondrial genes, *ND4*, *ND2*, *ATP6*, *CO1*, *TC*, and *TL1*. Remarkably, we noted that breaking the 5’-T rule did not obligately preclude mtDNA editing with DdCBEs. However, αDdCBEs consistently outperformed canonical DdCBEs, thereby supporting unconstrained mtDNA editing as a potential strategy for disease modeling and gene therapy applications.

## RESULTS

### Mitochondrial base editing with αDdCBEs

Given the ability of the modified TALE NT-αN to recognize all 5’ bases,^52^ we hypothesized that αDdCBEs can edit mtDNA as efficiently as standard, 5’-T-compliant DdCBEs, regardless of the TALE 5’-T rule. Moreover, we expected 5’-T-noncompliant DdCBEs to induce poor editing efficiencies relative to 5’-T-compliant DdCBEs. To test these hypotheses, we compared pairs of DdCBEs and αDdCBEs in both 5’-T-compliant and 5’-T-noncompliant formats at four mitochondrial loci in HEK293T cells (**Fig. 1**). To avoid variations in base editing outcomes as a result of spacer variability, we maintained fixed spacer sequences at each target locus by lengthening or shortening the TALE binding sites from the 5’ ends by a maximum of 2 bp each.

**Figure 1.**
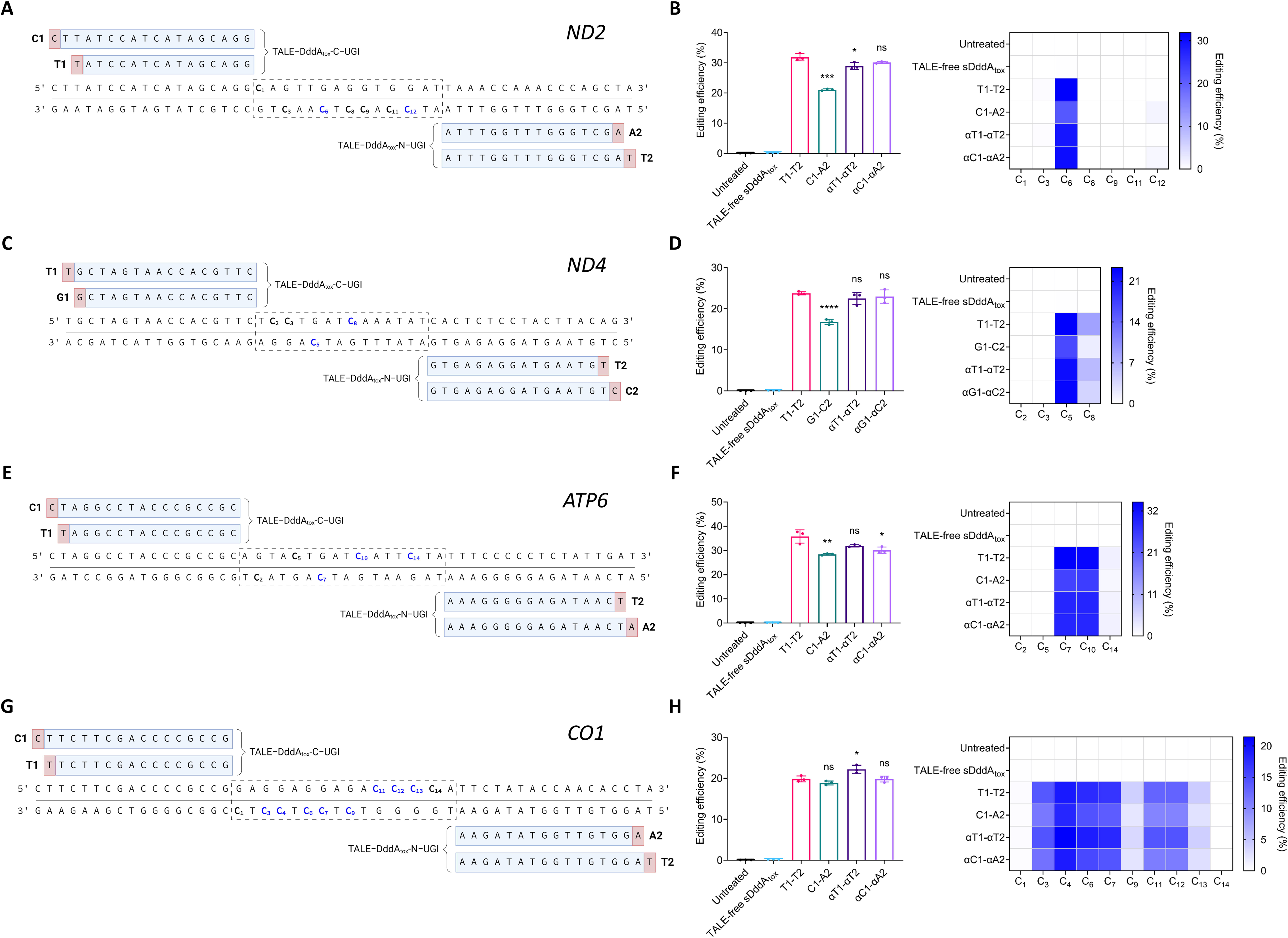
Mitochondrial base editing with 5’-T-compliant and 5’-T-noncompliant DdCBEs and αDdCBEs. **(A), (C), (E), (G)** Target sites within *ND2*, *ND4*, *ATP6*, and *CO1*, respectively. The sequences targeted by the TALE repeat arrays are shown in the blue rectangles, and the nucleotides immediately upstream of these sequences are indicated in the red boxes. Each DdCBE arm is denominated as N1/N2 (where ‘N’ represents A, C, G, or T) depending on its corresponding TALE 5’-N nucleotide and whether it constitutes the left (‘1’) or the right (‘2’) arm of the base editor. The spacers (sequences between TALE target sequences, where base editing is expected) are enclosed in dashed rectangles. All cytosines within the spacers are in bold and numbered relative to their positions from the 3’ end of their respective T1 TALE target sequences. Cytosines consistently edited at ≥1% across experimental conditions are highlighted in blue. **(B)**, **(D)**, **(F)**, **(H)** Overall editing efficiencies (left) and corresponding editing patterns (right) induced by 5’-T-compliant and 5’-T-noncompliand DdCBEs and αDdCBEs at *ND2*, *ND4*, *ATP6*, and *CO1*, respectively. TALE-free sDddA_tox_: N- and C-termini of TALE-free, mitochondrially targeted, split DddA_tox_–UGI. T1-T2: 5’-T-compliant DdCBE pairs. V1-V2 (where ‘V’ represents a non-T nucleotide): 5’-T-noncompliant DdCBE pairs. αT1-αT2: 5’-T-compliant αDdCBE pairs. αV1-αV2: 5’-T-noncompliant αDdCBE pairs. All measurements were obtained via NGS and correspond to editing efficiencies in HEK293T cells 3 days post-transfection. Values and error bars in **(B)**, **(D)**, **(F)**, and **(H)** represent the mean ± s.d. of *n* = 3 independent biological replicates. Displayed statistical significances were determined by comparing against the T1-T2 condition. \**P*<0.05; \*\**P*<0.01; \*\*\**P*<0.001; \*\*\*\**P*<0.0001; ns (not significant), *P*>0.05 by two-tailed unpaired *t* test in GraphPad Prism 10.

In addition to the untreated condition, TALE-free MTS–split DddA_tox_–UGI (labeled as TALE-free sDddA_tox_) was used as a negative control. Additionally, DdCBE pairs are generally referred to as N1-N2 (N = A, C, G, or T), denoting the 5’-most base of either the left (1) or the right (2) TALE. Accordingly, αDdCBEs are designated as αN1-αN2. Hence, T1-T2 denotes a 5’-T-compliant DdCBE pair, which corresponds to a positive control. Besides, unless otherwise noted, all base editors were designed with G1397-split DddA_tox_ in the C-to-N configuration, i.e., left TALE–DddA_tox_-C–UGI + right TALE–DddA_tox_-N– UGI.^13^

At the *ND2* locus, we observed that C1-A2 was the least active base editor, reaching overall editing efficiencies of ∼21%, whereas T1-T2, αT1-αT2, and αC1-αA2 displayed editing frequencies ranging from ∼29% to ∼32%. Additionally, all *ND2* base editors resulted in similar mutation patterns (**Fig. 1A, B**). Similarly, at the *ND4* site, we found that G1-C2 induced editing efficiencies of ∼17%, the lowest compared to T1-T2, αT1-αT2, and αG1-αC2, which installed edits at frequencies between ∼22% and ∼24%. Interestingly, both G1-C2 and αG1-αC2 resulted in more specific mutation patterns within the spacer than their 5’-T-compliant counterparts (**Fig. 1C, D**). Comparably, at the *ATP6* locus, C1-A2 was less efficient than T1-T2, with editing frequencies of ∼28% compared to ∼36%, respectively. Moreover, both αT1-αT2 and αC1-αA2, which displayed editing frequencies of up to ∼32%, were nearly as effective as T1-T2. Notably, all *ATP6* base editors displayed similar editing patterns (**Fig. 1E, F**). Unexpectedly, at the *CO1* site, all base editors displayed similar levels of activity and mutation patterns, with overall efficiencies ranging from ∼19 to ∼22%. Besides, despite the preference of DddA_tox_ for cytosines in TC motifs,^13^ several non-TC motifs within the *CO1* spacer were efficiently edited (**Fig. 1G, H**).

Collectively, these results suggest that canonical DdCBEs can effectively edit mtDNA even if their respective TALEs break the 5’-T rule. However, αDdCBEs tend to perform similarly to 5’-T-compliant DdCBEs and outperform 5’-T-noncompliant DdCBEs, thereby surpassing canonical DdCBEs in regard to design flexibility.

### Characterizing off-target editing by αDdCBEs

Seeking to characterize the specificity profiles of αDdCBEs relative to DdCBEs, based on an approach reported by Willis et al.,^29^ we calculated the normalized ratios between the on-target (i.e., within the spacer) and average amplicon-wide off-target editing efficiencies for each base editor. These quantities enabled us to conduct direct comparisons between the overall performance of αDdCBEs in contrast to DdCBEs, both in terms of their on-target editing activities and proximal off-target effects.

At the *ND2* locus, C1-A2, αT1-αT2, and αC1-αA2 resulted in an ∼1.8-fold reduction in average amplicon-wide off-target editing compared to T1-T2 (**Fig. 2A**). Accordingly, given their high on-target editing activities (**Fig. 1B**) and relatively low off-target effects (**Fig. 2B**, left), αDdCBEs considerably outperformed their canonical counterparts at the *ND2* site (**Fig. 2B**, right). In contrast, all *ND4* base editors introduced off-target editing at frequencies below 0.2% throughout the amplicon (**Fig. 2C**). However, given the relatively low on-target activity displayed by G1-C2 (**Fig. 1D**), and the moderately higher off-target effects caused by T1-T2 compared to the other pairs (**Fig. 2D**, left), at the *ND4* locus, αDdCBEs outperformed DdCBEs (**Fig. 2D**, right). Similarly, all *ATP6* base editors introduced off-target cytosine conversions at rates below 0.5% (**Fig. 2E**). However, both T1-T2 and C1-A2 resulted in somewhat higher average amplicon-wide off-target editing than αT1-αT2 and αC1-αA2 (**Fig. 2F**, left). Consequently, at the *ATP6* site, αDdCBEs performed better than DdCBEs (**Fig. 2F**, right). In contrast, at *CO1*, the 5’-T-compliant pairs resulted in higher average amplicon-wide off-target editing efficiencies compared to the 5’-T-noncompliant pairs (**Fig. 2G** and **Fig. 2H**, left). Therefore, both 5’-T-noncompliant *CO1* base editors outperformed the 5’-T-compliant DdCBE pair, T1-T2, with αC1-αA2 showing the highest overall performance (**Fig. 2H**, right).

**Figure 2.**
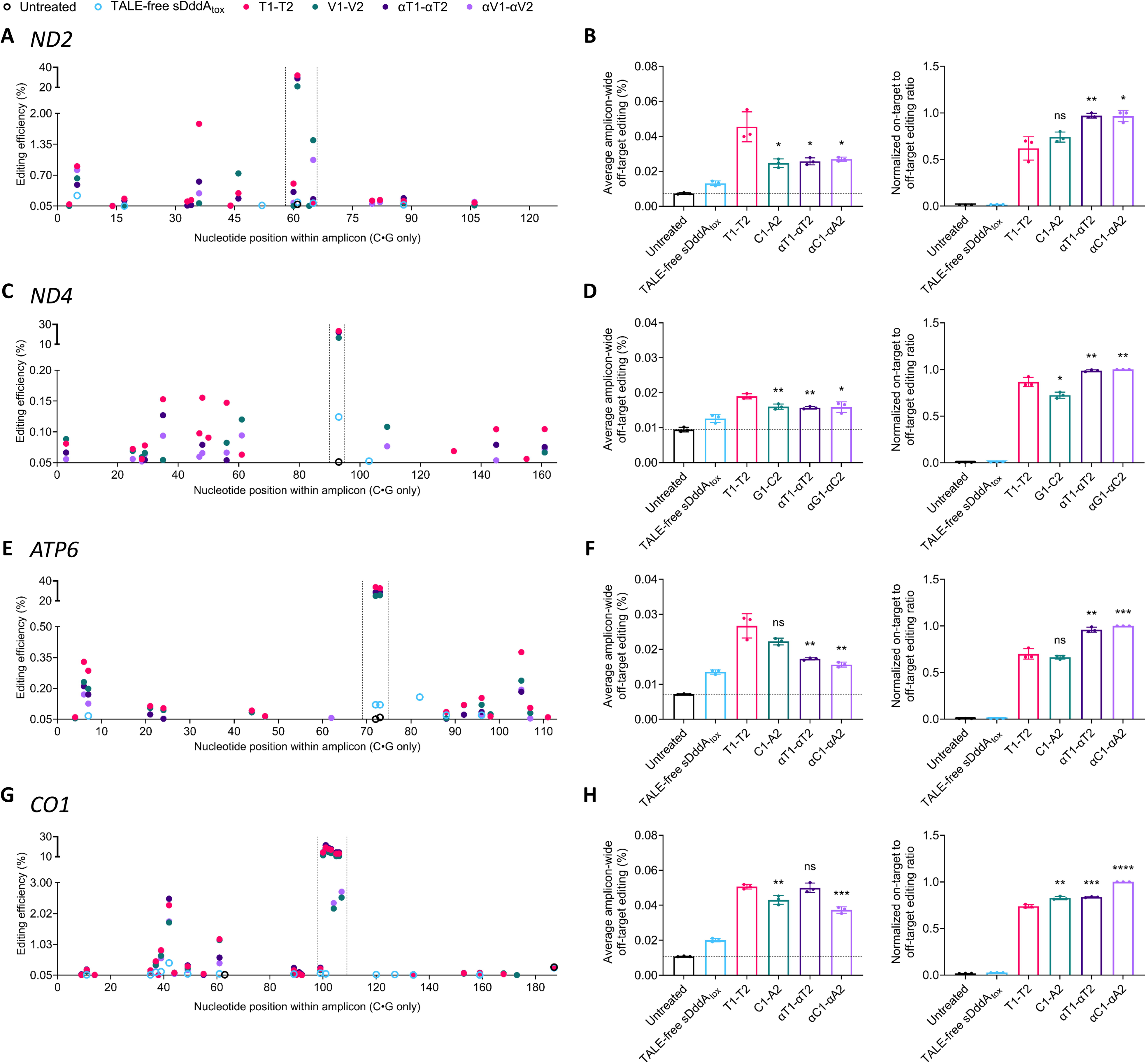
Proximal off-target effects by 5’-T-compliant and 5’-T-noncompliant DdCBEs and αDdCBEs. **(A)**, **(C)**, **(E)**, **(G)** Amplicon-wide editing efficiencies induced by 5’-T-compliant and 5’-T-noncompliant DdCBEs and αDdCBEs at *ND2*, *ND4*, *ATP6*, and *CO1*, respectively. The narrow regions between the vertical dashed lines confine the cytosines within the spacers at each target site. Note that the vertical axes are divided into two segments, with the ranges of the bottom segments adjusted to showcase the amplitudes of the proximal off-target effects, and the ranges of the top segments adjusted to showcase the amplitudes of the on-target editing events. All measurements were obtained via NGS and correspond to editing efficiencies in HEK293T cells 3 days post-transfection. All values represent the mean of *n* = 3 independent biological replicates. TALE-free sDddA_tox_: N- and C-termini of TALE-free, mitochondrially targeted, split DddA_tox_–UGI constructs. T1-T2: 5’-T-compliant DdCBE pairs. V1-V2 (where ‘V’ represents a non-T nucleotide): 5’-T-noncompliant DdCBE pairs. αT1-αT2: 5’-T-compliant αDdCBE pairs. αV1-αV2: 5’-T-noncompliant αDdCBE pairs. **(B)**, **(D)**, **(F)**, **(H)** Average amplicon-wide off-target editing (left) and normalized on-target to off-target editing ratios (right) of 5’-T-compliant and 5’-T-noncompliand DdCBEs and αDdCBEs at *ND2*, *ND4*, *ATP6*, and *CO1*, respectively. The horizontal dashed line in the average amplicon-wide off-target editing bar graphs corresponds to the mean of the untreated condition. Values and error bars in **(B)**, **(D)**, **(F)**, and **(H)** represent the mean ± s.d. of *n* = 3 independent biological replicates. Displayed statistical significances were determined by comparing against the T1-T2 condition. \**P*<0.05; \*\**P*<0.01; \*\*\**P*<0.001; \*\*\*\**P*<0.0001; ns (not significant), *P*>0.05 by two-tailed unpaired *t* test in GraphPad Prism 10.

Subsequently, to further characterize the specificity profiles of αDdCBEs relative to DdCBEs, we investigated their nuclear off-target effects at a TALE-dependent site (*MTND4P12*) and a TALE-independent site (chr8:37153384C, hg38) in *ND4*-edited cells (**Supplementary Fig. S1**).^13,53^ Notably, the *MTND4P12* off-target and the *ND4* on-target sequences differ by a single G/A mismatch (**Supplementary Fig. S1A**). Remarkably, at *MTND4P12*, T1-T2 resulted in off-target editing efficiencies of ∼16%. In contrast, all other pairs achieved frequencies of ∼0.2% (G1-C2), ∼7.5% (αT1-αT2), and ∼1% (αG1-αC2) (**Supplementary Fig. S1B,C**). On the other hand, at the herein examined TALE-independent off-target region, which was previously identified by Lei et al.^53^ as a frequently observed nuclear off-target across DdCBEs, and shares no sequence homology with the *ND4* on-target sequence, the editing efficiencies remained substantially similar among base editors. Nonetheless, αG1-αC2 resulted in moderately higher cytosine conversion rates relative to all other editors, albeit at efficiencies below 0.2%. Besides, TALE-free sDddA_tox_ led to off-target editing with frequencies of ∼1% (**Supplementary Fig. S1D**).

As a whole, these results suggest that, in the scope of proximal off-target effects in mtDNA and, potentially, nuclear editing at TALE-dependent off-target sites, αDdCBEs tend to be more specific than standard, 5’-T-compliant DdCBEs, thereby outperforming them in terms of specificity.

### Evaluating the generality of αDdCBE-induced mitochondrial base editing enhancements

We then explored whether αDdCBEs consistently led to base editing enhancements relative to DdCBEs, regardless of the 5’-most bases of their TALE binding sites. To this end, we identified *ATP6* as a locus accessible by base editors containing TALEs targeting sequences preceded by A, C, G, or T, with moderate variability across the resulting spacers. We designed TALE proteins containing between 15 and 17 repeats, and delimiting spacers ranging from 13 to 18 bp long. Of note, all spacers contained a bottom- and a top-strand cytosine approximately halfway through, and an additional top-strand cytosine towards the 3’ end (**Fig. 3A**).

**Figure 3.**
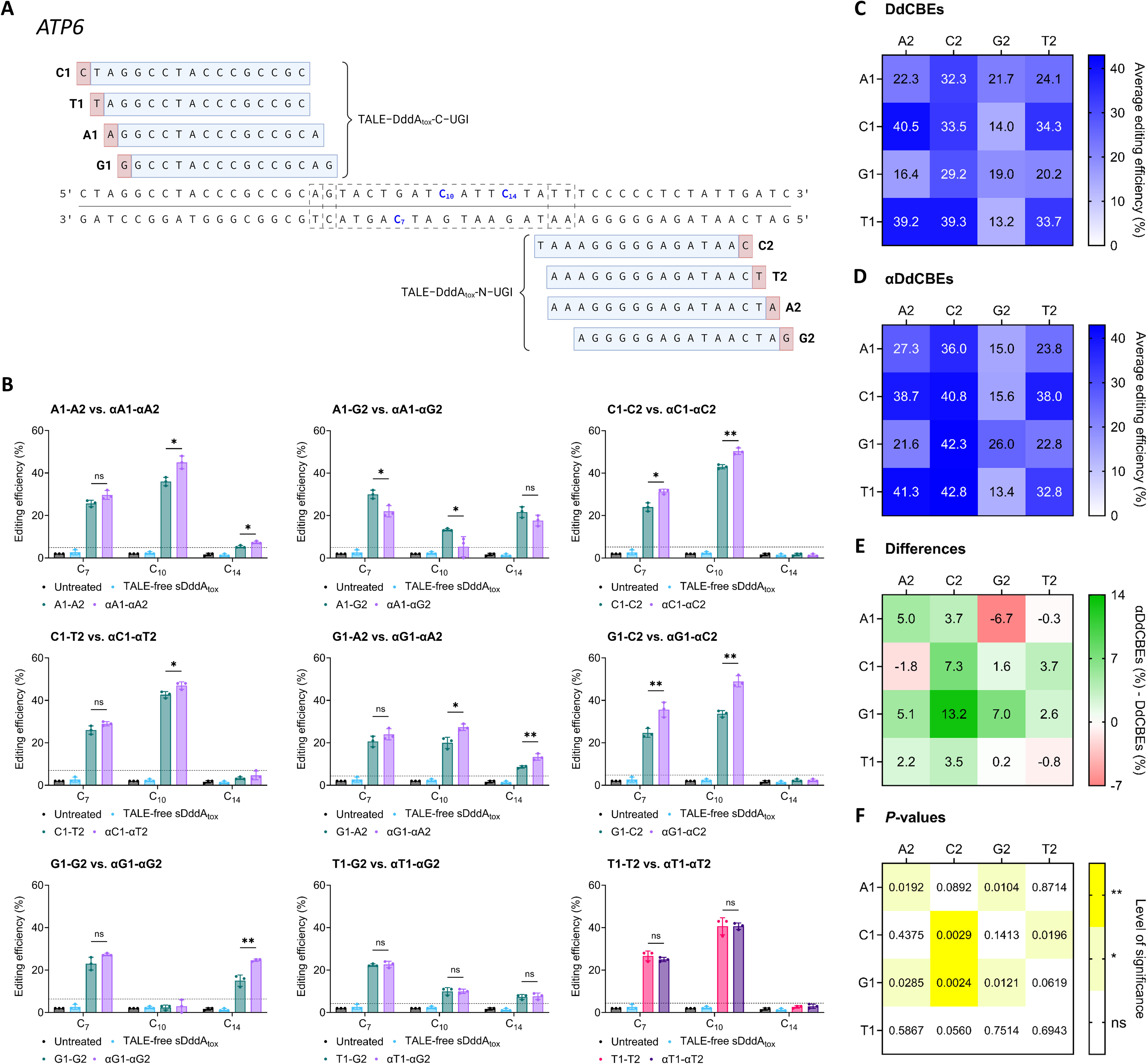
Evaluating αDdCBE-induced mitochondrial base editing enhancements at *ATP6*. **(A)** Target site within *ATP6*. The spacers are enclosed in dashed rectangles. Only the cytosines that were consistently edited across conditions are highlighted in bold blue and numbered from the 3’ end of the T1 TALE target sequence. **(B)** Representative comparisons between the editing efficiencies induced by DdCBE and αDdCBE pairs at *ATP6*. TALE-free sDddA_tox_: N- and C-termini of TALE-free, mitochondrially targeted, split DddA_tox_–UGI constructs. N1-N2 (where ‘N’ represents A, C, G, or T): canonical DdCBE pairs. αN1-αN2: αDdCBE pairs. T1-T2 and αT1-αT2 are shown in red and purple, respectively, to maintain the color scheme used in previous figures. All measurements were obtained via Sanger sequencing and correspond to editing efficiencies in HEK293T cells 3 days post-transfection. Values and error bars represent the mean ± s.d. of *n* = 3 independent biological replicates. The horizontal dashed lines correspond to critical percent values, obtained from sequencing trace decomposition analyses with a *P*-value cutoff of 0.01, above which base editing estimates are significantly different from background. **(C)**, **(D)** Average DdCBE- and αDdCBE-induced editing efficiencies at each specified N1-N2 condition. **(E)** Differences between the average editing efficiencies induced by αDdCBEs and DdCBEs. **(F)** Corresponding *P*-values for statistical comparisons between the average editing efficiencies displayed in **(C)** and **(D)**. \**P*<0.05; \*\**P*<0.01; ns (not significant), *P*>0.05 by two-tailed unpaired *t* test in GraphPad Prism 10.

We observed that equivalent base editors (i.e., A1-A2 and αA1-αA2, A1-C2 and αA1-αC2, and so on) led to similar mutation patterns, albeit frequently at moderately different efficiencies.

Representative comparisons between equivalent pairs are shown in **Fig. 3B**, and the complete set is displayed in **Supplementary Fig. S2**. Afterwards, we calculated the average activities of each construct to elucidate the general differences between αDdCBE- and DdCBE-induced base editing frequencies at *ATP6* (**Fig. 3C, D**). Then, we determined the differences between these quantities for each pair of equivalent base editors, along with their respective significance levels, facilitating the visualization of the overall αDdCBE-induced activity enhancements across comparisons (**Fig. 3E, F**). Notably, broad improvements in the efficiency of base editing with αDdCBEs relative to DdCBEs were observed in 6 out of the 16 total comparisons (**Fig. 3C–F**).

It is worth noting that *ATP6* base editors containing G2/αG2 arms were generally the least effective across conditions (**Fig. 3C, D**); moreover, only αA1-αG2 led to a statistically significant overall activity reduction relative to its canonical counterpart (**Fig. 3E, F**). Furthermore, only G2/αG2-containing pairs, except for G1-G2 and αG1-αG2, targeted spacer sequences with lengths of 17 or 18 bp (**Fig. 3A**). DddA-derived base editors targeting spacers of such lengths often install lower editing efficiencies compared to pairs with spacers up to 16 bp long.^13,54^ Thus, we hypothesized that reducing the spacers of G2/αG2-containing pairs to 16 bp or less would improve their editing efficiencies.

To test this hypothesis and characterize the effects of decreasing spacer length on the activities of A1-G2 vs. αA1-αG2, we evaluated two additional sets of arms: G2.16/αG2.16 and G2.17/αG2.17 (**Supplementary Fig. S3A**). Interestingly, G2.16/αG2.16-containing pairs led to lower editing efficiencies relative to the initial G2/αG2-containing pairs (designated as G2.15/αG2.15 in **Supplementary Fig. S3**), as well as minimal αDdCBE-induced enhancements. Strikingly, G2.17/αG2.17-containing pairs, with shorter spacers, led to improvements in base editing efficiencies relative to the original pairs, as well as greater αDdCBE-induced base editing reductions (**Supplementary Fig. S3B**).

Overall, these results further indicate that DdCBEs and αDdCBEs can effectively edit mtDNA in 5’-T-noncompliant formats. However, αDdCBEs can often lead to greater mtDNA editing efficiencies than their canonical counterparts, although in particular contexts the opposite can be observed.

### αDdCBEs outperform DdCBEs at mtDNA sites with stretches without 5’-T nucleotides

Subsequently, we evaluated the effectiveness of αDdCBEs at target sites that, based on standard design principles,^13^ cannot be accessed without breaking the TALE 5’-T rule. To this purpose, we first assessed an array of base editors at the tRNA-Cys-encoding gene *TC*. This locus is reportedly editable by mDdCBEs but not by dimeric DdCBEs.^48^ In detail, we tested a standard pair (A1-T2), two partially modified pairs (αA1-T2 and A1-αT2), and a fully modified pair (αA1-αT2). Additionally, we included two monomeric controls: mA1 and mT2, analogous to the left and right arms of the dimeric constructs (**Fig. 4A**). Unexpectedly, the dimeric base editors induced overall editing efficiencies ranging from ∼49% to ∼56%, well above mA1 (∼24%) and mT2 (∼14%) (**Fig. 4B**). Furthermore, A1-T2, αA1-T2, A1-αT2, and αA1-αT2 were considerably more specific than mA1 and mT2, which installed edits outside of the intended target sequence at frequencies of up to ∼6% and ∼3%, in that order (**Fig. 4C** and **Fig. 4D**, left). Therefore, the dimeric base editors far outperformed their monomeric counterparts, with αA1-αT2 exhibiting the greatest overall performance (**Fig. 4D**, right).

**Figure 4.**
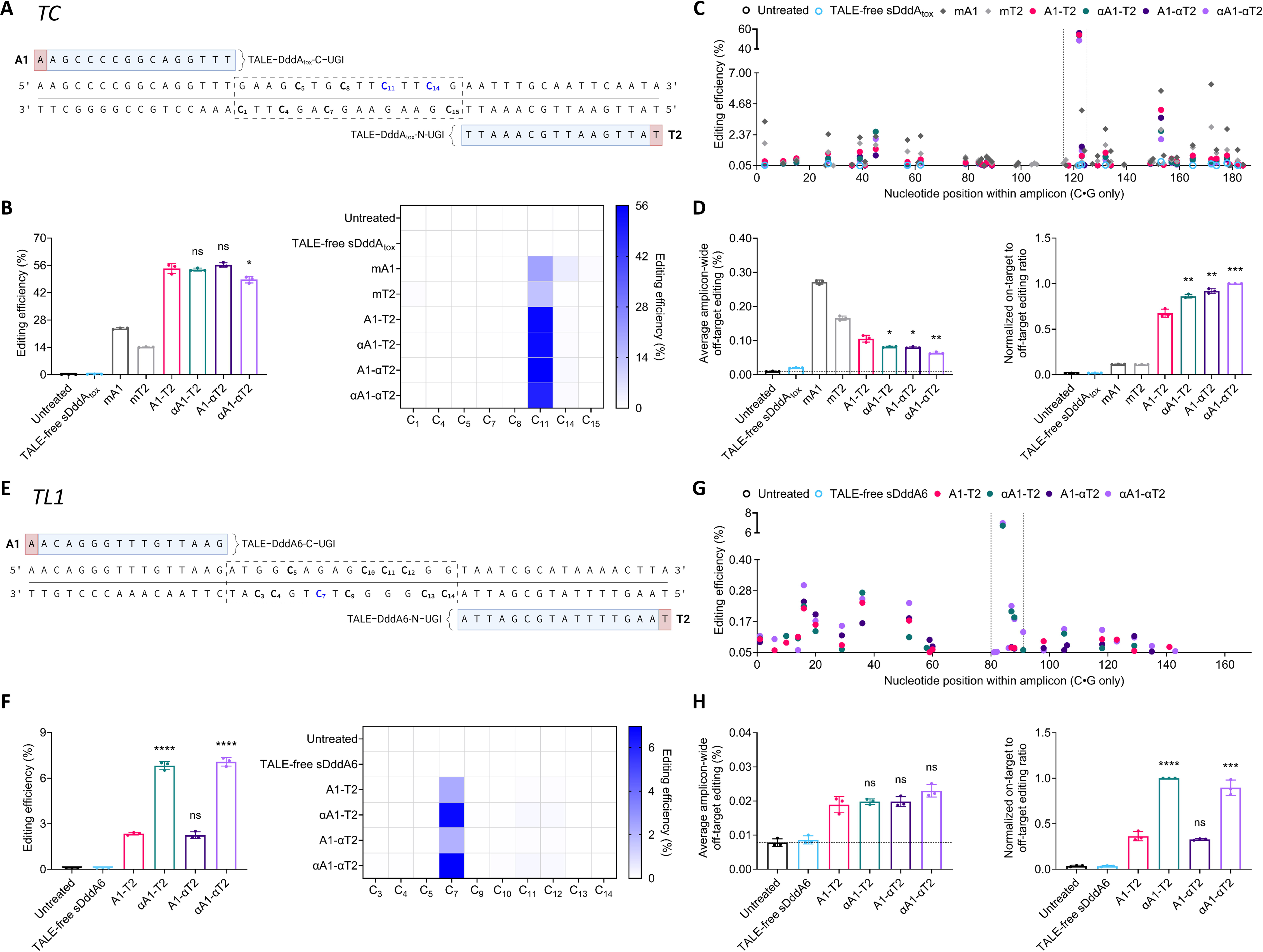
αDdCBEs outperform DdCBEs at mtDNA sites with stretches without 5’-T nucleotides. **(A), (E)** Target sites within the mitochondrial tRNA-Cys-encoding gene *TC* and the mitochondrial tRNA-Leu-encoding gene *TL1*, respectively. **(B), (F)** Overall editing efficiencies (left) and corresponding editing patterns (right). **(C), (G)** Amplicon-wide editing efficiencies. The narrow regions between the vertical dashed lines confine the on-target cytosines within the spacers. **(D), (H)** Average amplicon-wide off-target editing (left) and normalized on-target to off-target editing ratios (right). The horizontal dashed line in the average amplicon-wide off-target editing bar graphs corresponds to the mean of the untreated condition. TALE-free sDddA_tox_/sDddA6: N- and C-termini of TALE-free, mitochondrially targeted, split DddA_tox_/sDddA6–UGI constructs. mA1: 5’-T-noncompliant monomeric DdCBE (mDdCBE) control. mT2: 5’-T-compliant mDdCBE control. A1-T2: canonical DdCBE pair. αA1-T2 and A1-αT2: partially modified pairs. αA1-αT2: αDdCBE pair. All measurements were obtained via NGS and correspond to editing efficiencies in HEK293T cells 3 days post-transfection. Values and error bars in **(B)**–**(D)** and **(F)**–**(H)** represent the mean ± s.d. of *n* = 3 independent biological replicates. Displayed statistical significances were determined by comparing against the A1-T2 conditions. \**P*<0.05; \*\**P*<0.01; \*\*\**P*<0.001; \*\*\*\**P*<0.0001; ns (not significant), *P*>0.05 by two-tailed unpaired *t* test in GraphPad Prism 10.

To further evaluate the effectiveness of αDdCBEs at target sites lacking accessible 5’-Ts, we assessed a set of base editors at the tRNA-Leu-encoding gene *TL1*. Pathogenic variants in this gene, such as the broadly prevalent m.3243A>G, are linked to impaired oxidative phosphorylation and a wide range of complex disease outcomes.^55–57^ We initially observed that DddA_tox_ was poorly active at *TL1*; hence, to obtain an ample range of activities for comparison purposes, we used DddA6, an enhanced variant of DddA_tox_, which showed improved editing efficiencies at this site (**Supplementary Fig. S4**). Of note, the denominations of the *TL1* base editors are similar to those of the *TC* pairs (**Fig. 4E**). Besides, given that monomeric variants for DddA6 have yet to developed,^48,54^ *TL1*-specific monomeric controls were not included. Remarkably, the αA1-containing pairs led to an ∼3-fold increase in activity relative to A1-T2 and A1-αT2 (**Fig. 4F**). Moreover, all base editors resulted in similar specificity profiles; thus, the αA1-containing pairs significantly outperformed their A1-containing counterparts (**Fig. 4H**).

These results collectively suggest that dimeric DddA-derived base editors containing either canonical or unconstrained TALE NTDs, or both, can effectively access loci with stretches without 5’-Ts. Nevertheless, in some contexts, utilizing unconstrained TALEs to target sequences preceded by non-T nucleotides can facilitate the introduction of targeted modifications in mtDNA.

### Fine-tuning mitochondrial base editing outcomes via TALE shifting with αDdCBEs

The outcomes of DddA-derived base editors are partly determined by spacer length and the positions of the target cytosines within the spacer.^13,28,29,54^ Given that these determinants are contributed by the DNA-binding domains, targeting a particular locus with different pairs of TALEs can lead to diverse editing outcomes.^32,58^ Indeed, depending on their TALE proteins, the *ATP6* base editors developed in this study resulted in distinct mutation patterns (**Fig. 3B**). Based on these observations and the overall greater design flexibility offered by αDdCBEs compared to DdCBEs, we aimed to explore TALE shifting with αDdCBEs as a strategy to fine-tune mitochondrial base editing outcomes. To this end, we leveraged the herein optimized TALE formats for *TL1* editing (**Fig. 4E–H**). In detail, we tested a total of eight partially or fully modified base editors. For clarity, the former are denoted as α_L_DdCBEs, since only the left (L) arms contain unconstrained TALEs, and the latter are simply referred to as αDdCBEs (**Fig. 5A**).

**Figure 5.**
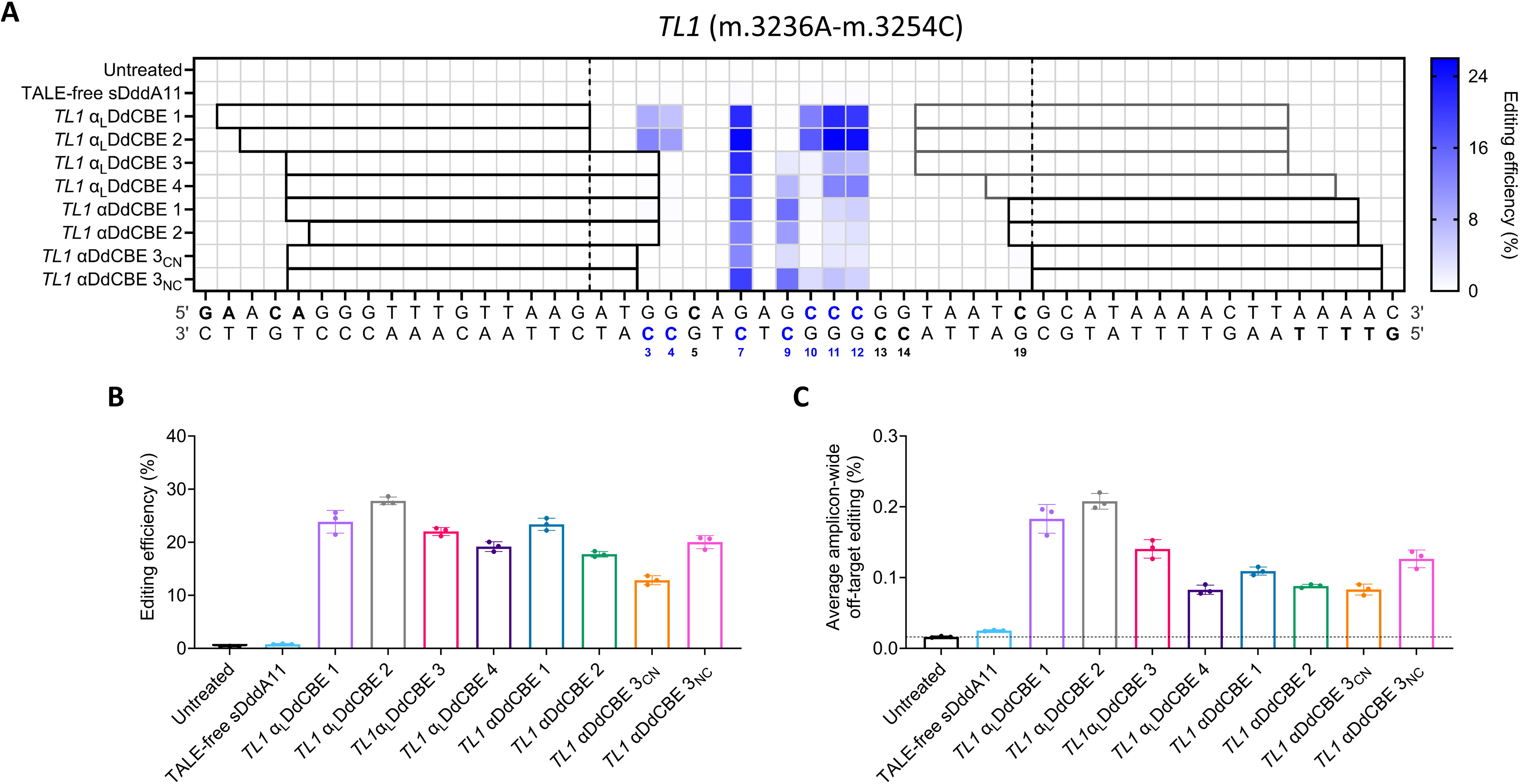
TALE shifting with αDdCBEs as a strategy to fine-tune mitochondrial base editing outcomes. **(A)** Heatmap detailing the editing patterns induced by DddA11-containing αDdCBEs and α_L_DdCBEs at *TL1*. The vertical dashed lines define an overall spacer region (m.3236A-m.3254C) by highlighting the 3’ end of the leftmost TALE target sequence and the 5’ end of the rightmost TALE target sequence. The 5’-most nucleotides of the TALE binding sites are shown in bold in the sequence below the heatmap. Cytosines within the overall spacer region are in bold and numbered relative to their positions from the left vertical dashed line. Cytosines that were edited at ≥1% are highlighted in blue. **(B)** Overall editing efficiencies induced by each base editor within their respective spacers. **(C)** Average amplicon-wide off-target editing efficiencies. The horizontal dashed line corresponds to the mean of the untreated condition. All measurements were obtained via NGS and correspond to editing efficiencies in HEK293T cells 3 days post-transfection. All values and error bars represent the mean ± s.d. of *n* = 3 independent biological replicates.

We presumed that promoting base editing across all cytosines within the *TL1* spacers would facilitate discerning the contributions of TALE shifting to mutation pattern variability. Thus, given that 9 out of the 11 cytosines within the *TL1* spacers are in non-T<u>C</u> motifs, which tend to be poorly processed by DddA_tox_ or DddA6,^13,54^ the base editors illustrated in **Fig. 5A** contain DddA11, a DddA_tox_ variant engineered to induce enhanced editing at T<u>C</u> and non-T<u>C</u> motifs,^54^ providing an adequate range of base editing outcomes for comparison purposes. Of note, *TL1* αDdCBE 3_NC_ corresponds to a pair with G1397-split DddA11 in the N-to-C configuration. All other pairs, including *TL1* αDdCBE 3_CN_, contain G1397-split DddA11 in the C-to-N orientation.

Broadly, we observed overall editing efficiencies ranging from ∼13% to ∼28% (**Fig. 5B**). Interestingly, *TL1* αDdCBEs generally displayed moderately improved levels of specificity compared to *TL1* α_L_DdCBEs (**Fig. 5C**). Regarding mutation patterns, C_7_ was efficiently edited by all pairs; besides, *TL1* α_L_DdCBEs 1 and 2, which displayed promiscuous editing profiles, were the only pairs to edit C_3_ and C_4_. In contrast, more than half the activity of *TL1* α_L_DdCBE 3 and αDdCBE 3_CN_ corresponded to editing at C_7_ (**Fig. 5A**). Furthermore, despite differing in spacer length by just 1 bp relative to *TL1* αDdCBEs 1 and 2, *TL1* α_L_DdCBE 4 displayed increased activity at C_11_ and C_12_, and decreased editing at C_9_. Moreover, likely due to their opposite orientations of split DddA11,^54^ *TL1* αDdCBEs 3_CN_ and 3_NC_ resulted in distinct mutation patterns.

In the context of DddA-derived base editors, these results collectively demonstrate that TALE shifting, i.e., utilizing different sets of TALEs to access a single target site, can result in enhanced editing outcomes, both in terms of activity and cytosine selectivity within the spacer regions.

## DISCUSSION

In this study, we formally developed αDdCBEs for unconstrained mitochondrial base editing, which display an expanded targeting scope, enhanced design flexibility, and improved specificity profiles compared to canonical DdCBEs. In addition, by functionally exploring the widely accepted TALE 5’-T rule for the design of effective DdCBEs, we found that this rule often acts as a moderate limiting factor for efficient mtDNA editing, rather than as an obligate constraint. Furthermore, we demonstrated that αDdCBEs are compatible with DddA_tox_ and its engineered variants, DddA6 and DddA11. Finally, we validated TALE shifting with αDdCBEs as an approach to fine-tune base editing outcomes. This method enables the definition of diverse spacers at a single target site, regardless of the 5’-most nucleotides available to the TALEs. In terms of practical applications, TALE shifting can be leveraged to modulate bystander editing.

Given the near universally excellent high activity of αDdCBEs at nearly all tested loci, we recommend their general use in mtDNA editing applications, focusing on spacer composition rather than TALE binding sites. Alternatively, if canonical DdCBEs result in suboptimal editing efficiencies and mutation patterns, αDdCBEs can be used to optimize these outcomes. When using αDdCBEs, we suggest defining spacers ranging from 11 to 18 bp long (preferably up to 16 bp), delimited by TALEs containing 15 or 16 repeats, irrespective of the 5’-most nucleotides of their target sequences. Regarding deaminase domain selection, our work is consistent with the previous recommendations by Mok et al.^54^ Additionally, we hypothesize that other effector domains, such as DddA homologs or chimeric deaminases,^16,32–36^ will also work well with unconstrained TALEs.

In this study, αDdCBEs were developed using immortalized human cells in vitro. Further characterizations in clinically relevant cell types and in vivo models are needed to continue to validate our observations. Likewise, although amplicon-wide analyses are highly informative,^29^ genome-wide surveys will enhance our understanding of the overall specificity of αDdCBEs.

In conclusion, we successfully developed efficient, specific, and sequence unconstrained mitochondrial base editors. These enhanced editors are compatible with various deaminase domains and can access virtually any sequence in human mtDNA. This design flexibility will facilitate their implementation in discovery science, disease modeling and gene therapy applications.

## MATERIALS AND METHODS

### Construction of FusX-compatible DdCBE backbone plasmids

All backbone plasmids were made via restriction cloning. A list including the source material for each construct is provided in **Supplementary Table S1**. In general, insert and vector bands were separated by agarose gel electrophoresis and purified with the Monarch DNA Gel Extraction Kit obtained from New England Biolabs (NEB). Ligations were done with the Quick Ligation^TM^ Kit (NEB). NEB^®^ Stable Competent *E. coli* (C3040H) were used for propagation, following the high efficiency transformation protocol specified by the manufacturer, and incubating plates and liquid cultures at 30 °C. Plasmids were purified with the QIAprep Spin Miniprep Kit (Qiagen) and sequence-verified via whole-plasmid sequencing (Primordium Labs).

### Assembly of DdCBE-encoding plasmids

All DdCBE-encoding plasmids used in this study were assembled via the FusX TALE Base Editor (FusXTBE) platform.^49–51^ Briefly, following standard design rules,^13,54^ DdCBEs were designed in silico with TALE Writer^50,51^ and SnapGene. Specifically, TALE repeat arrays were designed to target between 15 and 17 bp, separated by spacers ranging from 11 to 18 bp long.^13,50,54^ A list of all TALE binding sites is provided in **Supplementary Table S2**. DdCBE-encoding plasmids were assembled via Golden Gate cloning.^49–51^ Primers used for colony PCR (see **Supplementary Table S3**) were synthesized as standard DNA oligos by Integrated DNA Technologies (IDT). NEB^®^ Stable Competent *E. coli* (C3040H) were used for propagation. Plasmids were purified with the QIAprep Spin Miniprep Kit (Qiagen) and sequence-verified via whole-plasmid sequencing (Primordium Labs).

### Generation of TALE-free constructs

Plasmids encoding TALE-free MTS–G1397-split DddA_tox_/DddA6/DddA11–UGI were generated with the Q5^®^ Site-Directed Mutagenesis (SDM) Kit (NEB), using the corresponding herein developed backbone plasmids as templates for each final construct containing either the DddA_tox_, DddA6, or DddA11 C- or N-terminal halves.^13,54^ Cloning was carried out following the manufacturer’s instructions with the provided NEB^®^ 5-alpha Competent *E. coli* cells. Primers for SDM (see **Supplementary Table S3**) were designed using NEBaseChanger (NEB) and synthesized as standard DNA oligos (IDT). Plasmids were purified with the QIAprep Spin Miniprep Kit (Qiagen) and sequence-verified via whole-plasmid sequencing (Primordium Labs).

### Mammalian cell culture and lipofection

HEK293T cells (CRL-3216^TM^, ATTC) were maintained at 37 °C and 5% CO_2_. The cells were cultured in high glucose DMEM (Thermo Fisher Scientific) supplemented with 10% (v/v) fetal bovine serum (Thermo Fisher Scientific) and 100 U ml^-1^ penicillin-streptomycin (Thermo Fisher Scientific). Lipofectamine^TM^ 3000 Transfection Reagent (Thermo Fisher Scientific) was used for lipofections. In brief, 24 h before lipofection, 0.3×10^6^ cells/well were seeded in 6-well plates. Then, lipofections proceeded with 500 ng per monomer for DdCBEs and TALE-free constructs to make up 1,000 ng of total plasmid DNA,^13,54^ and 1,000 ng of plasmid DNA for monomeric DdCBEs (mDdCBEs).^48^ Cells were collected for genotyping at 72 h post-transfection.

### Genomic DNA isolation from mammalian cell culture

At 72 h post-transfection, cell medium was aspirated, the cells were washed with 500 µl 1x DPBS without calcium or magnesium (Thermo Fisher Scientific), trypsinized with 500 µl 1x Trypsin-EDTA (0.5%) without phenol red (Thermo Fisher Scientific) for 5 minutes at 37 °C and collected in microcentrifuge. Total genomic DNA (including mitochondrial DNA) was purified using the DNeasy Blood & Tissue Kit (Qiagen) following the manufacturer’s instructions and stored at -20 °C until further downstream processing.

### High-throughput sequencing of genomic DNA samples

Genotyping primers were designed using Primer-BLAST,^59^ querying the Homo sapiens genome assembly hg38 for primer pair specificity. In detail, to increase PCR specificity for the intended mitochondrial target sequences, PCR was biased against the amplification of nuclear mitochondrial pseudogenes (NUMTs)^60^ by aligning the 3’ ends of candidate primers with specific single-nucleotide mismatches between intended mitochondrial targets and potential unintended nuclear templates, and accordingly adding 3’-terminal phosphorothioate (PS) bonds to the primers. This strategy avoids 3’-terminal editing of the mismatched primers by the 3’-5’ exonuclease activity of Q5^®^ High-Fidelity DNA Polymerase (NEB), increasing PCR specificity.^61^

Primers including the partial Illumina^®^ forward and reverse adapter sequences, in addition to barcodes for sample multiplexing, were synthesized as Ultramer^TM^ DNA oligos (IDT). Afterward, genomic sites of interest were amplified with the Q5^®^ High-Fidelity 2X Master Mix (NEB) using conventional thermocycling conditions. Then, PCR products corresponding to the same experimental condition but with different barcodes were combined after agarose gel electrophoresis, purified using the Monarch DNA Gel Extraction Kit (NEB), confirmed via Sanger sequencing (Genewiz), and submitted to NGS (Amplicon-EZ with partial adapters, Genewiz). Alternatively, if the samples were not multiplexed, the PCR products were individually purified with the QIAquick PCR Purification Kit (QIAGEN), confirmed via Sanger sequencing (Genewiz), and submitted to NGS (Amplicon-EZ without partial adapters, Genewiz). A list of all genotyping primers is provided in **Supplementary Table S4**.

### Analysis of high-throughput sequencing data

In multiplexed samples, the paired-end read FASTQ files generated by NGS were demultiplexed and analyzed utilizing the CRISPRessoPooled tool within CRISPResso2.^62,63^ Similarly, if the samples were not multiplexed, the paired-end read FASTQ files were analyzed with the CRISPRessoBatch tool within CRISPResso2.^62,63^ In general, DdCBE spacer sequences were used as the guide sequence input. Besides, for each replicate in each experimental condition, the sequence of the amplicon corresponding to the target site, plus the respective barcode if demultiplexing, was used as the amplicon sequence input. Unless otherwise stated, the quantification window size was set to 8 or 10, and the quantification window center was set to -8 or -10. All optional parameters were set to NA.^13,62^

The output alleles frequency table was used to determine the overall on-target editing in each sample, calculated as the percentage of aligned reads with C•G-to-T•A conversions within a spacer.^48^ Likewise, the output nucleotide percentage table was used to calculate the editing activity at each cytosine within each spacer, as well as the proximal off-target editing within each amplicon.^29,48^ In detail, similar to the methodology followed in the development of zinc-finger DdCBEs (ZF-DdCBEs), average amplicon-wide off-target editing was quantified as the sum of all C•G-to-T•A conversions within an amplicon, excluding its corresponding DdCBE spacer, over the total number of C•G base pairs within that amplicon.^29^ Calculations were done in Microsoft Excel.

### Targeted amplicon sequencing for nuclear DNA off-target analyses

Based on previous reports, nested PCR was performed to amplify a TALE-dependent off-target site within the NUMT *MTND4P12*, and conventional PCR to amplify a frequent TALE-dependent off-target site at chr8:37153384C (hg38).^13,53,64^ Primers for the first PCR (PCR1) in the nested PCR strategy were synthesized as standard DNA oligos (IDT). Primers for the generation of amplicons for NGS were synthesized as Ultramer^TM^ DNA oligos (IDT), including barcodes and Illumina^®^ adapters. PCR was done with the Q5^®^ High-Fidelity 2X Master Mix (NEB). After PCR1 in the nested PCR strategy, amplicons were purified with the QIAquick PCR Purification Kit (Qiagen), and 10 ng were used as template DNA for the second PCR. Amplicons for targeted deep sequencing were purified as detailed in “High-throughput sequencing of genomic DNA samples” and submitted to NGS (Amplicon-EZ with partial adapters, Genewiz).

The methodology for data analysis was similar to the approach described in “Analysis of high-throughput sequencing data.” In detail, to determine the overall nuclear DNA off-target editing at *MTND4P12*, the CRISPResso2 output alleles frequency table was used to calculate the percentage of aligned reads with C•G-to-T•A conversions within the pseudo-spacer (i.e., the nuclear DNA region analogous to the genuine target spacer in mtDNA). Additionally, the output nucleotide percentage table was used to calculate the editing activity at each cytosine within the pseudo-spacer. Similarly, the nucleotide percentage table was used to quantify the nuclear DNA off-target editing at the above-mentioned TALE-independent off-target locus.

### Sanger sequencing of genomic DNA samples and data analysis

For the 5’ nucleotide precedence analyses at the *ATP6* locus, genomic DNA from *ATP6*-edited cells and the corresponding controls was purified as described in “Genomic DNA isolation from mammalian cell culture”. Afterward, PCR was conducted with the Q5^®^ 2X Master Mix (NEB) and *ATP6* Sanger sequencing primers (listed in **Supplementary Table S4**), which were designed as detailed in “High-throughput sequencing of genomic DNA samples” (without adapters or barcodes) and synthesized as standard DNA oligos with 3’ PS bonds (IDT). Then, PCR products were visualized by electrophoresis in a 1% agarose gel, purified with the QIAquick PCR Purification Kit (QIAGEN), and submitted to Sanger sequencing (Genewiz).

The resulting trace (.ab1) files were analyzed in the EditR server.^65^ Briefly, DdCBE spacer sequences were used as the guide sequence input.^50^ Additionally, the 5’ starts and 3’ ends of the trace files were trimmed to exclude bases with quality scores lower than 40, and the *P*-value cutoff for calling base editing was set to 0.01. Besides, to exclude noise from low-confidence measurements in the calculation of the average editing efficiencies, these were computed per replicate as the mean of the predicted editing at cytosines within the spacer that corresponded to highly significant (*P* ≤ 0.01) editing events, as compared to untreated controls. Statistical analyses were conducted using two-tailed unpaired *t* tests in GraphPad Prism 10.

## ACKNOWLEDGMENTS

We thank Dr. David Liu, Dr. Beverly Mok, and colleagues (Broad Institute of MIT and Harvard) for their work on the development of DdCBEs and for kindly providing various DdCBE plasmids.

## DATA AVAILABILITY STATEMENT

FusX-compatible DdCBE backbone plasmids, as well as canonical and unconstrained DdCBE plasmids will be made available through Addgene. The FusX assembly system is available on Addgene (Kit # 1000000063). Source data for the main figures are provided in the supplementary material.

## AUTHOR CONTRIBUTION STATEMENT

**Santiago R. Castillo:** Conceptualization, Methodology, Software, Validation, Formal Analysis, Investigation, Data Curation, Writing – Original Draft, Writing – Review & Editing, Visualization, Project Administration, Funding Acquisition. **Brandon W. Simone:** Methodology, Writing – Review & Editing. **Karl J. Clark:** Writing – Review & Editing, Funding Acquisition. **Patricia Devaux:** Resources, Writing – Review & Editing, Supervision. **Stephen C. Ekker:** Resources, Writing – Review & Editing, Supervision, Project Administration, Funding Acquisition.

