## Supplementary Material for "Unconstrained Precision Mitochondrial Genome Editing with αDdCBEs"

**Page**

**Supplementary Figure S1.** Nuclear off-target effects induced by DdCBEs and αDdCBEs 2

**Supplementary Figure S2.** All N1-N2 vs. αN1-αN2 comparisons at *ATP6* 3

**Supplementary Figure S3.** Additional N1-G2 vs. αN1-αG2 comparisons at *ATP6* 5

**Supplementary Figure S4.** $DddA_{tox}$- vs. DddA6-containing *TL1* $\alpha_L$DdCBEs 7

**Supplementary Table S1.** Summary of subcloning strategies to generate all backbone plasmids 8

**Supplementary Table S2.** TALE binding sites of all base editors used in this study 9

**Supplementary Table S3.** Primers for cloning TALE-free constructs and colony PCR 10

**Supplementary Table S4.** Primers for Sanger sequencing and NGS 11

**Supplementary Sequences.** Architecture of base editors used in this study 13

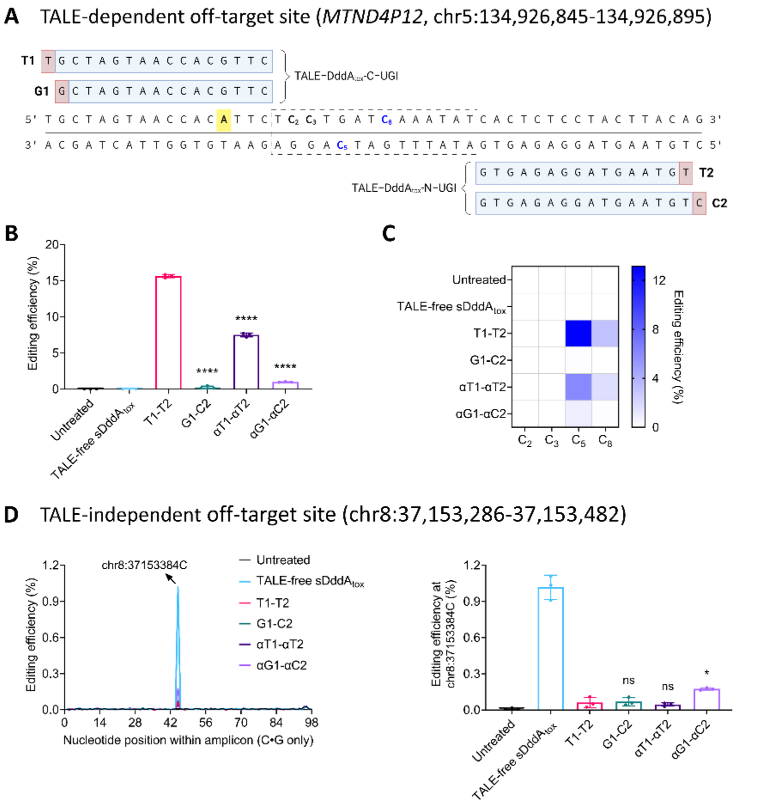


**Supplementary Figure S1. Nuclear off-targets effects induced by DdCBEs and αDdCBEs. (A)** TALE-dependent off-target site at the nuclear mitochondrial pseudogene *MTND4P12*, which differs from the mitochondrial sequence targeted by the indicated *ND4* base editors by a single nucleotide mismatch (highlighted in yellow).[1,2] Cytosines in the pseudo-spacer (dashed box) are numbered from the 3' end of the left TALE off-target sequence. Edited cytosines are highlighted in blue. **(B)** Overall off-target editing efficiencies and **(C)** corresponding mutation patterns at *MTND4P12*. **(D)** Left, amplicon-wide visualization of the editing efficiencies at a TALE-independent off-target site, which shares no sequence similarity with the *ND4* DdCBE on-target sequence.[2] Right, off-target editing at chr8:37153384C (hg38). TALE-free $sDddA_{tox}$: N- and C-termini of TALE-free, mitochondrially targeted, split $DddA_{tox}$–UGI constructs. T1-T2: 5'-T-compliant DdCBE pair; G1-C2: 5'-T-noncompliant DdCBE pair; αT1-αT2: 5'-T-compliant αDdCBE pair; αG1-αC2: 5'-T-noncompliant αDCBE pair. All pairs correspond to *ND4*-specific base editors. All measurements were obtained via NGS and correspond to editing efficiencies in HEK293T cells 3 days post-transfection. Values and error bars in **(B)** through **(D)** represent the mean ± s.d. of $n = 3$ independent biological replicates. Displayed statistical significances in **(B)** and **(D)** were determined by comparing against the T1-T2 condition. *$P$<0.05; ****$P$<0.0001; ns (not significant), $P$>0.05 by two-tailed unpaired *t* test in GraphPad Prism 10.

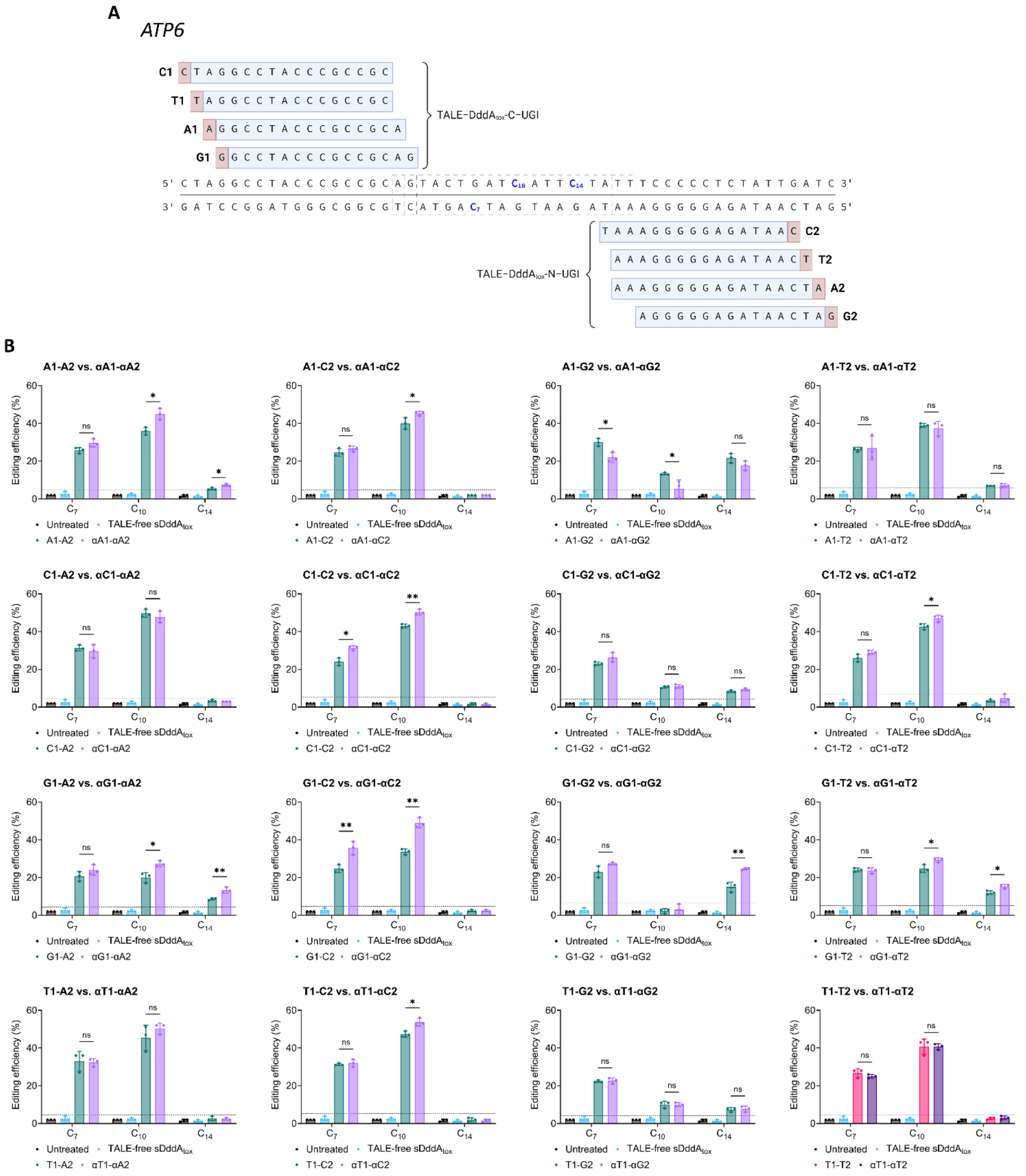


**Supplementary Figure S2. All N1-N2 vs. αN1-αN2 comparisons at *ATP6*. (A)** Schematic of the mitochondrial on-target site within *ATP6*. The spacer regions are indicated by the dashed boxes. Cytosines that were consistently edited across conditions are numbered and highlighted in blue from the 3' end of the T1 TALE target sequence. **(B)** Comparisons between the editing efficiencies induced by

DdCBEs and αDdCBEs in every possible N1-N2 combination, from A1-A2 (top left) to T1-T2 (bottom right, shown in red and purple to maintain the color scheme used in other figures). TALE-free sDddA$_{tox}$: N- and C-termini of TALE-free, mitochondrially targeted, split DddA$_{tox}$–UGI constructs. All measurements were obtained via Sanger sequencing trace decomposition with EditR[3] and correspond to editing efficiencies in HEK293T cells 3 days post-transfection. Values and error bars represent the mean ± s.d. of $n$ = 3 independent biological replicates. The horizontal dashed lines correspond to critical percent values, obtained from EditR[3] with a $P$-value cutoff of 0.01, above which base editing estimates are significantly different from background. *$P$<0.05; **$P$<0.01; ns (not significant), $P$>0.05 by two-tailed unpaired $t$ test in GraphPad Prism 10.

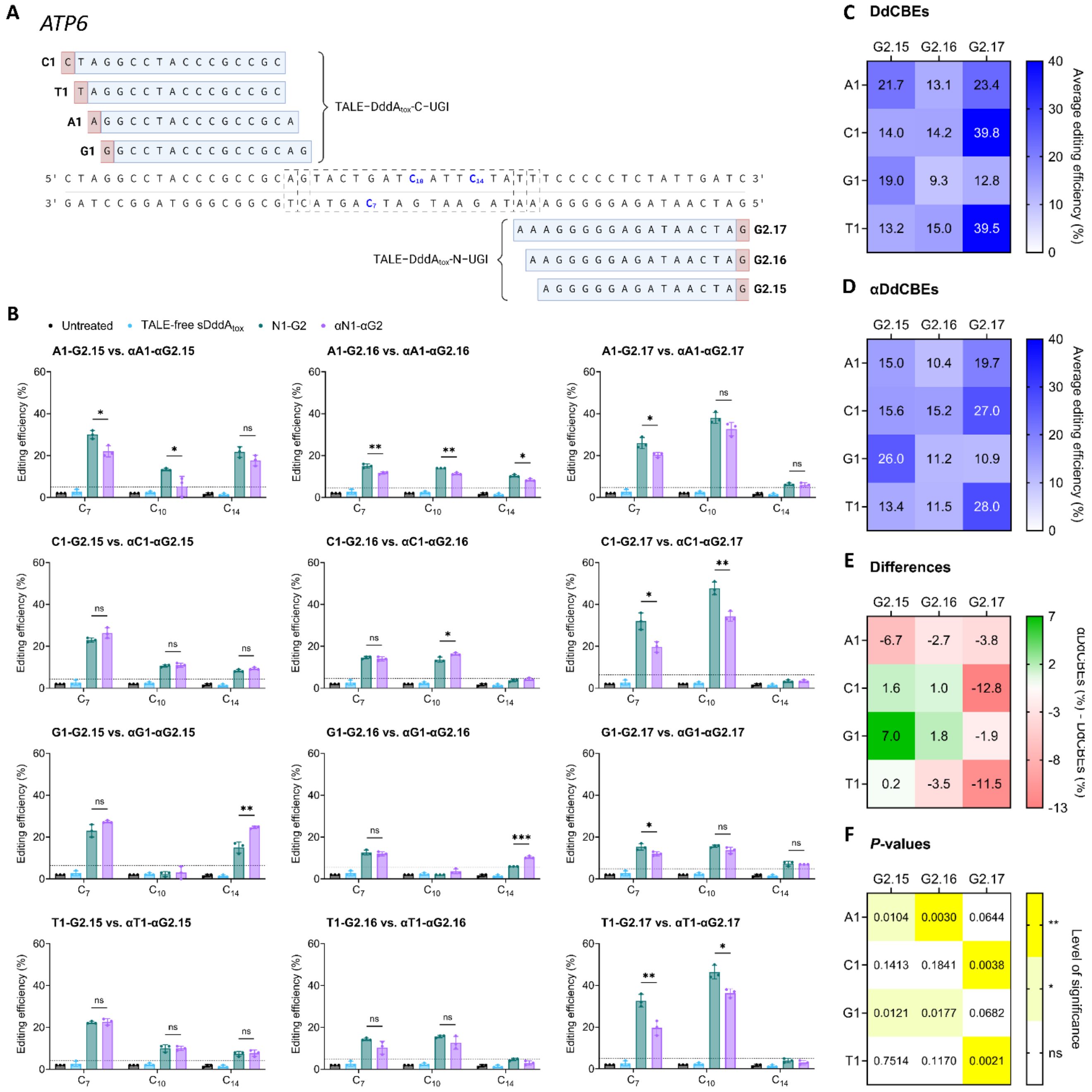


**Supplementary Figure S3. Additional N1-G2 vs. αN1-αG2 comparisons at *ATP6*. (A)** Schematic of the mitochondrial on-target site within *ATP6*. The spacer regions are indicated by the dashed boxes. Cytosines that were consistently edited across conditions are numbered and highlighted in blue from the 3' end of the T1 TALE target sequence. G2.15, which targets a 15 bp long sequence preceded by a guanine, corresponds to the right DdCBE arm used in the main comparisons in **Fig. 3** and **Supplementary Fig. S2**. Similarly, G2.16 and G2.17 correspond to right DdCBE arms that target 16 and 17 bp long sequences, respectively, both preceded by the same guanine. **(B)** Comparisons between the editing efficiencies induced by DdCBEs and αDdCBEs in every possible N1-G2 combination. All measurements were obtained via Sanger sequencing trace decomposition with EditR[3] and correspond to editing efficiencies in HEK293T cells 3 days post-transfection. Values and error bars represent the mean ± s.d. of $n$ = 3 independent biological replicates. The horizontal dashed lines correspond to critical percent values, obtained from EditR[3] with a $P$-value cutoff of 0.01, above which base editing estimates are significantly different from background. TALE-free sDddA$_{tox}$: N- and C-termini of TALE-free, mitochondrially targeted, split DddA$_{tox}$–UGI constructs. **(C)**, **(D)** Average DdCBE- and αDdCBE-induced editing efficiencies. **(E)** Differences between the average editing efficiencies induced by αDdCBEs and DdCBEs. **(F)** Corresponding $P$-values for statistical comparisons between the average editing efficiencies displayed in **(C)** and **(D)**. *$P$<0.05; **$P$<0.01; ns (not significant), $P$>0.05 by two-tailed unpaired $t$ test in GraphPad Prism 10.

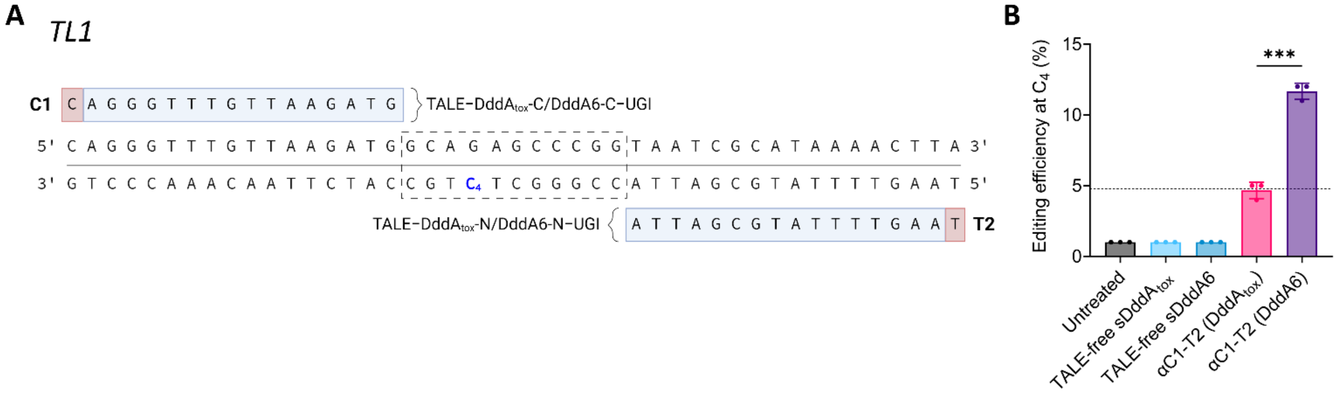


**Supplementary Figure S4. DddA$_{tox}$- vs. DddA6-containing *TL1* α$_L$DdCBEs. (A)** Schematic of the mitochondrial on-target site within *TL1* and preliminary α$_L$DdCBEs (the left arm contains an unconstrained TALE and the right arm a canonical TALE) tested to select an optimal effector domain. The spacer is indicated by the dashed box. $C_4$, highlighted in blue, which is numbered relative to its position from the 3’ end of the left TALE target sequence, corresponds to the only edited cytosine within the spacer. **(B)** Editing efficiencies at $C_4$ across conditions. TALE-free sDddA$_{tox}$/sDddA6: N- and C-termini of TALE-free, mitochondrially targeted, split DddA$_{tox}$/DddA6–UGI constructs. All measurements were obtained via Sanger sequencing trace decomposition with EditR[3] and correspond to editing efficiencies in HEK293T cells 3 days post-transfection. Values and error bars represent the mean ± s.d. of $n$ = 3 independent biological replicates. The horizontal dashed line corresponds to a critical percent value, obtained from EditR[3] with a $P$-value cutoff of 0.01, above which base editing estimates are significantly different from background.

**Supplementary Table S1. Summary of subcloning strategies to generate all backbone plasmids.** Vector and insert plasmids used to clone all FusX-compatible DdCBE backbones used in this study. Insert plasmid pUC57 FusX, containing a 'FusX cassette', was obtained through a gene synthesis service (GenScript). The FusX cassette consists of a fragment of the canonical TALE N-terminal domain (TALE NT-T), a BsmBI restriction site, the components necessary for blue-white screening (i.e., CAP binding site–lac promoter–lac operator–lacZα), an additional BsmBI restriction site, and a fragment of the TALE C-terminal domain. TALE NT-αN, unconstrained TALE N-terminal domain.[1,4–9]

| Backbone plasmid | Vector plasmid | | | Insert plasmid | | | Enzymes |
|---|---|---|---|---|---|---|---|
| | Source | Origin | Replaced | Source | Origin | Inserted | |
| DdCBE 1397C | ND5.1-DdCBE-right side TALE | Dr. David Liu (Broad Institute)[1] | ND5.1 right TALE | pUC57 FusX | This study | FusX cassette | PstI and BamHI |
| DdCBE 1397N | ND4-DdCBE-right side TALE | Addgene plasmid no. 157843[1] | ND4 right TALE | pUC57 FusX | This study | FusX cassette | PstI and BamHI |
| αDdCBE 1397C | ND6-DdCBE-right side TALE | Addgene plasmid no. 157841[1] | ND6 right TALE | pUC57 FusX | This study | FusX cassette | PstI and BamHI |
| αDdCBE 1397N | DdCBE 1397N | This study | TALE NT-T | αDdCBE 1397C | This study | TALE NT-αN | NheI and PstI |
| DdCBE DddA6/11 1397C | DdCBE 1397N | This study | DddAtox 1397C | ND5.2-Left DdCBE-G1397C-T1413I | Dr. David Liu (Broad Institute)[8] | DddAtox 1397C (T1413I) | BamHI and Bsu36I |
| DdCBE DddA6 1397N | ND5.2-Right TALE-G1397-N-DddA6 | Dr. David Liu (Broad Institute)[8] | ND5.2 right TALE | pUC57 FusX | This study | FusX cassette | PstI and BamHI |
| DdCBE DddA11 1397N | ND5.2-Right TALE-G1397-N-DddA11 | Dr. David Liu (Broad Institute)[8] | ND5.2 right TALE | pUC57 FusX | This study | FusX cassette | PstI and BamHI |
| αDdCBE DddA6/11 1397C | DdCBE DddA6/11 1397C | This study | TALE NT-T | αDdCBE 1397C | This study | TALE NT-αN | NheI and PstI |
| αDdCBE DddA6 1397N | DdCBE DddA6 1397N | This study | TALE NT-T | ND6-DdCBE-right side TALE | Addgene plasmid no. 157841[8] | TALE NT-αN | SacI and PstI |
| αDdCBE DddA11 1397N | αDdCBE 1397C | This study | DddA$_{tox}$ 1397C | ND5.2-Right TALE-G1397-N-DddA11 | Dr. David Liu (Broad Institute)[8] | DddA11 1397N | BamHI and Bsu36I |
| mDdCBE | DdCBE 1397N | This study | DddA$_{tox}$ 1397N | pCMV-ND1 Right-GSVG-UGI | Addgene plasmid no. 187413[9] | DddA$_{tox}$ GSVG | BamHI and Bsu36I |

**Supplementary Table S2. TALE binding sites of all base editors used in this study.** For simplicity, arms with the same TALE target sequence are grouped together. In the *TL1* section, α/$\alpha_L$-1/2/3/4 (left/right) refers to the base editors in **Fig. 5**. E.g., ‘$\alpha_L$-1 (left)’ corresponds to the left arm of *TL1* $\alpha_L$DdCBE 1. For each TALE, its corresponding 5’-N nucleotide is specified ($N_0$), as well as the target sequence of its central repeat domain (CRD). Also, the plasmids utilized for FusX-based assembly are listed on the right.

| Target | Base editor arm | TALE target seq. (5'-to-3') | | FusX plasmids for assembly[4–6] | | | | | | |
|---|---|---|---|---|---|---|---|---|---|---|
| | | $N_0$ | CRD target seq. | X1 | X2 | X3 | X4 | B2 | B3 | LR |
| *ATP6* | A1/αA1 | A | GGCCTACCCGCCGCA | 42 | 29 | 22 | 38 | 10 | - | NI |
| | A2/αA2 | A | TCAATAGAGGGGGAAA | 53 | 13 | 35 | 43 | - | 33 | NI |
| | C1/αC1 | C | TAGGCCTACCCGCCGC | 51 | 38 | 50 | 23 | - | 23 | HD |
| | C2/αC2 | C | AATAGAGGGGGAAAT | 4 | 9 | 43 | 41 | 1 | - | NG |
| | G1/αG1 | G | GCCTACCCGCCGCAG | 38 | 50 | 23 | 23 | 5 | - | NN |
| | G2.15/αG2.15 (G2/αG2) | G | ATCAATAGAGGGGGA | 14 | 4 | 9 | 43 | 11 | - | NI |
| | G2.16/αG2.16 | G | ATCAATAGAGGGGGAA | 14 | 4 | 9 | 43 | - | 41 | NI |
| | G2.17/αG2.17 | G | ATCAATAGAGGGGGAAA | 14 | 4 | 9 | 43 | - | 41 | 11 |
| | T1/αT1 | T | AGGCCTACCCGCCGC | 11 | 24 | 6 | 26 | 7 | - | HD |
| | T2/αT2 | T | CAATAGAGGGGGAAA | 17 | 51 | 11 | 43 | 1 | - | NI |
| *CO1* | A2/αA2 | A | GGTGTTGGTATAGAA | 44 | 48 | 44 | 13 | 9 | - | NI |
| | C1/αC1 | C | TTCTTCGACCCCGCCG | 62 | 62 | 34 | 22 | - | 38 | NN |
| | T1/αT1 | T | TCTTCGACCCCGCCG | 56 | 55 | 6 | 23 | 6 | - | NN |
| | T2/αT2 | T | AGGTGTTGGTATAGAA | 11 | 60 | 59 | 52 | - | 9 | NI |
| *ND2* | A2/αA2 | A | GCTGGGTTTGGTTTA | 40 | 43 | 64 | 44 | 16 | - | NI |
| | C1/αC1 | C | TTATCCATCATAGCAGG | 61 | 54 | 14 | 13 | - | 37 | 16 |
| | T1/αT1 | T | ATCCATCATAGCAGG | 14 | 20 | 20 | 10 | 3 | - | NN |
| | T2/αT2 | T | AGCTGGGTTTGGTTTA | 10 | 59 | 48 | 59 | - | 64 | NI |
| *ND4* | C2/αC2 | C | TGTAAGTAGGAGAGTG | 60 | 3 | 51 | 35 | - | 12 | NN |
| | G1/αG1 | G | CTAGTAACCACGTTC | 29 | 45 | 6 | 7 | 16 | - | HD |
| | T1/αT1 | T | GCTAGTAACCACGTTC | 40 | 12 | 2 | 18 | - | 48 | HD |
| | T2/αT2 | T | GTAAGTAGGAGAGTG | 45 | 12 | 11 | 9 | 12 | - | NN |
| *TC* | mA1/A1/αA1 | A | AGCCCCGGCAGGTTT | 10 | 22 | 42 | 11 | 16 | - | NG |
| | mT2/T2/αT2 | T | ATTGAATTGCAAATT | 16 | 33 | 63 | 17 | 4 | - | NG |
| *TL1* | A1/αA1/$\alpha_L$-2 (left) | A | ACAGGGTTTGTTAAG | 5 | 43 | 64 | 48 | 1 | - | NN |
| | αC1/$\alpha_L$-3/$\alpha_L$-4/α-1 (left) | C | AGGGTTTGTTAAGATG | 11 | 48 | 60 | 49 | - | 36 | NN |
| | T2/αT2/$\alpha_L$-1/$\alpha_L$-2/$\alpha_L$-3 (right) | T | AAGTTTTATGCGATTA | 3 | 64 | 52 | 39 | - | 16 | NI |
| | $\alpha_L$-1 (left) | G | AACAGGGTTTGTTAAG | 2 | 11 | 48 | 60 | - | 49 | NN |
| | $\alpha_L$-4 (right) | T | TTAAGTTTTATGCGA | 61 | 12 | 64 | 15 | 7 | - | NI |
| | α-1/α-2 (right) | T | TTTAAGTTTTATGCG | 64 | 3 | 64 | 52 | 10 | - | NN |
| | α-2 (left) | A | GGGTTTGTTAAGATG | 43 | 64 | 48 | 3 | 4 | - | NN |
| | α-$3_{CN}$/α-$3_{NC}$ (left) | C | AGGGTTTGTTAAGAT | 11 | 48 | 60 | 49 | 9 | - | NG |
| | α-$3_{CN}$/α-$3_{NC}$ (right) | G | TTTTAAGTTTTATGC | 64 | 49 | 48 | 61 | 15 | - | HD |

**Supplementary Table S3. Primers for cloning TALE-free constructs and colony PCR.** Plasmids encoding TALE-free, mitochondrially targeted split deaminase domain–UGI were generated via site-directed mutagenesis (SDM) with the primers indicated below, which were designed using NEBaseChanger (NEB) and synthesized as standard DNA oligos (IDT).

| Template | Primers | | TALE-free construct |
|---|---|---|---|
| | Name | Sequence (5'-to-3') | |
| DdCBE 1397N<br>DdCBE DddA6 1397N<br>DdCBE DddA11 1397N | DddAN F | GGATCCGGCAGCTACGCC | TALE-free DddA$_{tox}$-N–UGI<br>TALE-free DddA6-N–UGI<br>TALE-free DddA11-N–UGI |
| | 3xFLAG R1 | CATCTTGTCATCGTCATCCTTGTAATCGATG | |
| DdCBE 1397C<br>DdCBE DddA6/11 1397C | DddAC F | GGATCCGCCATTCCAGTG | TALE-free DddA$_{tox}$-C–UGI<br>TALE-free DddA6/11-C–UGI |
| | 3xFLAG R2 | CATCTTGTCATCGTCATCCTTG | |

Additionally, colony PCR in FusX-based assembly[6] was conducted with the standard DNA oligos (IDT) indicated below. For assemblies with backbone plasmids DdCBE DddA6 1397N or DdCBE DddA11 1397N, FusX F2 was used. For assemblies with all other backbone plasmids, FusX F1 was used.

| Primers | |
|---|---|
| Name | Sequence (5'-to-3') |
| FusX F1 | CTACCCATGAAGCGATTGTG |
| FusX F2 | CCACCCATGAAGCTATTGTG |
| FusX R | ATCCACCCAGGCCTTTCTTC |

### Supplementary Table S4. Primers for Sanger sequencing and NGS.

| Target | Primers | |
|---|---|---|
| | Name | Sequence (5'-3') |
| *ATP6* | *ATP6* F | caccataattacccccatact*c |
| | *ATP6* R | gtcctttagtgttgtgtatgg*t |
| | *ATP6* BC1 F | ACACTCTTTCCCTACACGACGCTCTTCCGATCT**ATCACG**caccataattacccccatact*c |
| | *ATP6* BC2 F | ACACTCTTTCCCTACACGACGCTCTTCCGATCT**CGATGT**caccataattacccccatact*c |
| | *ATP6* BC3 F | ACACTCTTTCCCTACACGACGCTCTTCCGATCT**TTAGGC**caccataattacccccatact*c |
| | *ATP6* NGS R | GACTGGAGTTCAGACGTGTGCTCTTCCGATCTgtcctttagtgttgtgtatgg*t |
| *CO1* | *CO1* NGS F | ACACTCTTTCCCTACACGACGCTCTTCCGATCTttctccttacacctagcaggt*g |
| | *CO1* BC1 R | GACTGGAGTTCAGACGTGTGCTCTTCCGATCT**ATCACG**gctcgtgtgtctacgtctatt |
| | *CO1* BC2 R | GACTGGAGTTCAGACGTGTGCTCTTCCGATCT**CGATGT**gctcgtgtgtctacgtctatt |
| | *CO1* BC3 R | GACTGGAGTTCAGACGTGTGCTCTTCCGATCT**TTAGGC**gctcgtgtgtctacgtctatt |
| *ND2* | *ND2* BC1 F | ACACTCTTTCCCTACACGACGCTCTTCCGATCT**ATCACG**ggttacccaaggcaccc |
| | *ND2* BC2 F | ACACTCTTTCCCTACACGACGCTCTTCCGATCT**CGATGT**ggttacccaaggcaccc |
| | *ND2* BC3 F | ACACTCTTTCCCTACACGACGCTCTTCCGATCT**TTAGGC**ggttacccaaggcaccc |
| | *ND2* NGS R | GACTGGAGTTCAGACGTGTGCTCTTCCGATCTggtgcgagatagtagtagggt*c |
| *ND4* | *ND4* BC1 F | ACACTCTTTCCCTACACGACGCTCTTCCGATCT**ATCACG**ttctcataatcgcccacgg*g |
| | *ND4* BC2 F | ACACTCTTTCCCTACACGACGCTCTTCCGATCT**CGATGT**ttctcataatcgcccacgg*g |
| | *ND4* BC3 F | ACACTCTTTCCCTACACGACGCTCTTCCGATCT**TTAGGC**ttctcataatcgcccacgg*g |
| | *ND4* NGS R | GACTGGAGTTCAGACGTGTGCTCTTCCGATCTggttgagggataggaggag*a |
| *TC* | *TC* NGS F | ACACTCTTTCCCTACACGACGCTCTTCCGATCTcctacctatctccccttttat*a |
| | *TC* BC1 R | GACTGGAGTTCAGACGTGTGCTCTTCCGATCT**ATCACG**ccaatgtctttgtggtttgtag |
| | *TC* BC2 R | GACTGGAGTTCAGACGTGTGCTCTTCCGATCT**CGATGT**ccaatgtctttgtggtttgtag |
| | *TC* BC3 R | GACTGGAGTTCAGACGTGTGCTCTTCCGATCT**TTAGGC**ccaatgtctttgtggtttgtag |
| *TL1* | *TL1* F | cgtttgttcaacgattaaag |
| | *TL1* R | agcgaagggttgtagtagcc |
| | *TL1* BC1 F | ACACTCTTTCCCTACACGACGCTCTTCCGATCT**ATCACG**cgtttgttcaacgattaaag |
| | *TL1* BC2 F | ACACTCTTTCCCTACACGACGCTCTTCCGATCT**CGATGT**cgtttgttcaacgattaaag |
| | *TL1* BC3 F | ACACTCTTTCCCTACACGACGCTCTTCCGATCT**TTAGGC**cgtttgttcaacgattaaag |
| | *TL1* NGS R | GACTGGAGTTCAGACGTGTGCTCTTCCGATCTagcgaagggttgtagtagcc |
| *MTND4P12* | TASd F1 | ggataggaggaggataggggata |
| | TASd R1 | cggcgcagtcattctcata*g |
| | TASd BC1 F2 | ACACTCTTTCCCTACACGACGCTCTTCCGATCT**ATCACG**ttaatgtggtggctgagcg |
| | TASd BC2 F2 | ACACTCTTTCCCTACACGACGCTCTTCCGATCT**CGATGT**ttaatgtggtggctgagcg |
| | TASd BC3 F2 | ACACTCTTTCCCTACACGACGCTCTTCCGATCT**TTAGGC**ttaatgtggtggctgagcg |
| | TASd NGS R2 | GACTGGAGTTCAGACGTGTGCTCTTCCGATCTcaagcctcactaatctcgcc |
| *chr8:37153286-37153482 (hg38)* | TASi BC1 F | ACACTCTTTCCCTACACGACGCTCTTCCGATCT**ATCACG**ggagaaacgtcacggtatgc |
| | TASi BC2 F | ACACTCTTTCCCTACACGACGCTCTTCCGATCT**CGATGT**ggagaaacgtcacggtatgc |
| | TASi BC3 F | ACACTCTTTCCCTACACGACGCTCTTCCGATCT**TTAGGC**ggagaaacgtcacggtatgc |
| | TASi NGS R | GACTGGAGTTCAGACGTGTGCTCTTCCGATCTggtggtcaacaattctcactgc |

Non-bold, capital letters in the primer sequences correspond to partial Illumina® adapter sequences. Bold, capital letters correspond to barcodes for multiplexing. Lower case letters correspond the annealing sites of the primers. Asterisks (*) denote phosphorothioate bonds.[10]

*ATP6* F and *ATP6* R were used to obtain the Sanger sequencing data in **Fig. 3**, **Supplementary Fig. S2**, and **Supplementary Fig. S3**. *TL1* F and *TL1* R were used to obtain the next-generation sequencing (NGS) data in **Fig. 5**, and the Sanger sequencing data in **Supplementary Fig. S4**. TASd F1 and TASd R1 were used for the first PCR in the nested PCR strategy to amplify the TALE-dependent off-target site within *MTND4P12* (**Supplementary Fig. S1**). All other primers were used to obtain the NGS data in the rest of the figures. All Sanger sequencing and NGS reactions were done through Genewiz.

**Supplementary Sequences S1. Architecture of base editors used in this study**

All DdCBE and αDdCBE arms, and monomeric DdCBEs (mDdCBEs) used in this study have the general architecture (from N- to C-terminus): COX8A MTS–3xFLAG–TALE NTD–TALE repeat array–TALE CTD–2aa linker–split deaminase domain or full-length DddA$_{tox}$ GSVG–4aa linker–UGI–ATP5B 3'UTR.

Below are the amino acid sequences of the two TALE N-terminal domains (TALE NT-T and TALE NT-αN) the general form of a FusX-based TALE repeat array, and the TALE C-terminal domain.[1,4,7] The differences between the TALE N-terminal domains are underlined in their respective sequences. The repeat variable diresidue (RVD) motifs are shown as XX in the TALE repeat array sequence. Similarly, the three dots (…) separate the first repeat from the last half repeat. All repeats in between have the same composition as the first repeat. The RVD-to-nucleotide correspondence is NI = A, HD = C, NN = G, and NG = T.[4]

**TALE NT-T**[1,7]

DIADLRTLGYSQQQQEKIKPKVRSTVAQHHEALVGHGFTHAHIVALSQHPAALGTVAVKYQDMIAALPEA
THEAIVGVGKQWSGARALEALLTVAGELRGPPLQLDTGQLLKIAKRGGVTAVEAVHAWRNALTGAPLN

**TALE NT-αN**[1,7]

DIADLRTLGYSQQQQEKIKPKVRSTVAQHHEALVGHGFTHAHIVALSQHPAALGTVAVKYQDMIAALPEA
THEAIVGVGKRGAGARALEALLTVAGELRGPPLQLDTGQLLKIAKRGGVTAVEAVHAWRNALTGAPLN

**FusX-based TALE repeat array**[4]

LTPDQVVAIASXXGGKQALETVQRLLPVLCQDHG…LTPDQVVAIASXXGGKQALE

**TALE C-terminal domain**[1]

SIVAQLSRPDPALAALTNDHLVALACLGGRPALDAVKKGLG

The amino acid sequences of the remaining components are available in their respective sources.[1,8,9]
